# Clock genes *Period 1* and *Period 2* in the hippocampal CA1 mediate depression-like behaviors and rapid antidepressant response

**DOI:** 10.1101/2021.08.14.456364

**Authors:** Xin-Ling Wang, Xiao-Xing Liu, Kai Yuan, Ying Han, Yan-Xue Xue, Shi-Qiu Meng, Su-Xia Li

## Abstract

Accumulated reports have indicated that circadian rhythm is closely related to the pathogenesis of major depressive disorder (MDD). Recently, adenosine has been identified to modulate circadian clock via adenosine A_1_ and A_2A_ receptor signaling pathways. Cyclic AMP-response element binding protein (CREB) is a convergent point that plays a critical role in the pathogenesis of depression and is a downstream molecule of adenosine A_1_ receptor signaling pathway as an endpoint that can regulate the expression of circadian genes *Period1* (*Per1*) and *Period2* (*Per2*). However, whether *Per* mediates the development of MDD via CREB has not been elucidated. We used chronic unpredictable stress (CUS) to induce depression-like behaviors and found that it could induce decrease in *p*-CREB and PER1 levels in the hippocampal CA1 region in rats. Both depression-like behaviors and the decreased protein levels could be rapidly rescued by the administration of adenosine A_1_ receptor agonist 2-Choro-N6-cyclopentyladenosine (CCPA). Furthermore, knockdown of *Per1* in hippocampal CA1 region could also induce depression-like behaviors, which could also be rescued by CCPA. Interestingly, *Per2* knockdown in hippocampal CA1 region resulted in potential antidepressant-like effect. In addition, knockout of CRE sequence in the promoter regions of either *Per1* or *Per2* led to depression-like behaviors, which could not be rescued by CCPA. These results indicated that clock genes *Per1* and *Per2* play critical roles in the pathophysiology of depression and CRE sequences in the promoter regions of *Per1* and *Per2* may be a critical antidepressant target.

**Highlights:**

1. CUS induces both depression-like behaviors and decreases in the expression of *p*-CREB and PER1 levels in the hippocampal CA1 region in rats, which can be rapidly rescued by 2-Choro-N6-cyclopentyladenosine (CCPA).
2. Knockdown of clock gene *Per1* in the hippocampal CA1 brain region leads to depression-like behaviors in rats, which can be also rescued by CCPA.
3. Knockdown of clock gene *Per2* in the hippocampal CA1 brain region may have potential antidepressant-like effect.
4. Knockout of the CRE sequence on the promoter region of the clock genes *Per1* and *Per2* produces depression-like behaviors, which cannot be rescued by CCPA.

## Introduction

Major depressive disorder (MDD) is a chronic relapsing psychiatric disease. The clinical symptoms are anhedonia, loss of interest and circadian rhythm disturbances. The most dangerous symptom is suicide. Multiple types of antidepressants have been put into clinical use, but most of them take at least two weeks to take effect, which limits the prevention of suicide in patients with MDD. Therefore, further study on the pathogenesis of depression and development of fast onset of action antidepressants has become a top priority.

Recent years, several reviews described that circadian disturbances are involved in the pathogenesis of MDD and the rapid antidepressant therapies [1–3]. This viewpoint is based on animal model studies to find abnormalities of the clock gene rhythmical expression, or through clinical researches to find closely relationship between clock gene polymorphism and the occurrence of depression. In addition, abnormal expression of clock genes in brain [4] and peripheral blood [5] has been revealed in patients with MDD. Orozco-solis et al. [6] analyzed common candidate genes and cell pathways in mice treated with sleep deprivation or ketamine through comparative genomics studies, and found that both treatments could cause down-regulated transcription of 5 clock genes (*Per2, Npas4, Rorb, Dbp, and Ciart*). However, these previous researches only revealed the correlation between the pathophysiology of MDD and biorhythm from phenomenology, there is no in-depth study in the molecular biology process.

Several studies have reported that adenosine A_1_ receptor (A_1_R) agonists (including 2-Choro-N6-cyclopentyladenosine (CCPA), MRS5474, etc.) have rapid antidepressant-like effects [7, 8]. MRS5474 showed its rapid antidepressant-like action within one hour after injection [7]. By contrast, CCPA had a sedative effect at the first few hours after injection, and showed its rapid antidepressant-like effect later in rodents [8]. However, the detailed mechanisms have been rarely reported [7]. Although it has been demonstrated that A_1_R agonist can regulate circadian genes [9–11], this pathophysiology has not been linked to depression at present.

Previous study showed that cyclic AMP response element-binding protein (CREB) was a downstream molecule of the A_1_R signaling pathway [12]. It has been well documented that CREB is a convergent molecule in the pathogenesis of MDD and in the signaling pathways of antidepressant therapies [13–15]. Phosphorylated CREB could translocate into nucleus and bind to the cAMP-responsive elements (CREs) in the promoter regions of numerous genes, including clock genes *Period 1* (*Per1*) and *Period 2* (*Per2*), which are core circadian genes [15–18]. Therefore, we hypothesized that clock gene *Per1* and *Per2* might play critical roles in the pathophysiology of MDD via CRE sequence in their promoter regions. In this study, we used CCPA as an activator of adenosine signaling to activate its downstream molecule of CREB, *Per1/2* gene knockdown technique in hippocampal CA1 in rats, and CRE sequence knockout strategy in the promoter region of *Per1/2* gene in rats to verify our hypothesis.

## Materials and Methods

### Animals

Male Sprague Dawley (SD) rats (220-240 g) were bought from the Laboratory Animal Center, Peking University Health Science Center and housed in groups of five under a reverse 12 h/12 h light/dark cycle (8:00 light off and 20:00 light on) with free access to food and water. The room temperature was maintained 23±1°C and humidity was 50±5% controlled. SD *Per1* ^(c.-3353_-3345delTGACGTCA) Smoc^ and SD *Per2* ^(c.-10368_-10360delTGACGTCA) Smoc^ rats were custom-made from Cyagen Company (Jiangsu, China). They were propagated, and the descendants were identified to select homozygotes in our lab to do the following experiments. All of the animal procedures were performed in accordance with the National Institutes of Health Guide for the Care and Use of Laboratory Animals and were approved by the Biomedical Ethics Committee for Animal Use and Protection of Peking University (Ethical approval number: LA2011-017 and LA2019063).

### Drugs

2-Choro-N6-cyclopentyladenosine (CCPA) (Abcam, Cambridge, UK) was dissolved in 2% dimethylsulfoxide and 98% saline before use. CCPA was given to rats intraperitoneally.

### Intracranial microinjections

After the rats were anaesthetized by isoflurane gas, we microinjected AAV-sh*Per1/*sh*Per2* (1ul per side) or AAV-Scramble (1ul per side) bilaterally into CA1 (anterior/posterior, −4.3 mm; medial/lateral, ±2.0 mm; and dorsal/ventral, −2.0 mm [19, 20] with Hamilton syringes connected to 30-gauge injectors. We injected within 5 minutes and kept the injector in place for an additional 3 minutes to allow diffusion.

### Tissue sample preparation and western blot

We based the procedures of western blot on our previous studies [19, 21]. Specially, membrane proteins were extracted by using membrane and cytoplasmic protein extraction kits (Bi Yun Tian, Shanghai, China).

### Sucrose preference test (SPT), forced swim test (FST), Open field test (OFT) and elevated plus maze (EPM)

The procedures were based on previous studies [22–24]. **See Supplementary Information for details**.

### Chronic unpredictable stress (CUS) protocol

CUS was performed with two different stressors per day for 28 days, which was adapted from previous studies [23, 25]. **See Supplementary Information for details**.

### Locomotor activity

Locomotor activity (LA) was detected with an automated video-tracking system (DigBehv-LM4; Shanghai Jiliang Software Technology, Shanghai, China), which had been used in our previous studies [26]. A monochrome video camera was mounted on top of each chamber. All of the chambers were connected to a computer. The video files (stored on the computer) were analyzed using DigBehv analysis software. LA was expressed as the total distance traveled during the 5 min test.

### Statistical analysis

The data were expressed as mean±SEM. Statistical analyses were performed using Prism 5 software (GraphPad). The statistical analyses of data were done using unpaired two-tailed Student’s *t*-test, one-way analysis of variance (ANOVA), two-way ANOVA followed by Bonferroni’s multiple tests as appropriate, which can be referred to the figure legends. Values of *P* < 0.05 were considered statistically significant.

## Results

### CUS induced depression- and anxiety-like behaviors and decreased *p*-CREB-PER1 signaling in hippocampal CA1

First, we verified that CCPA had a potential antidepressant-like effect using FST (Fig S1a, b). The decreased immobility time in the FST could persist from 24 h to 36 h after administration of CCPA (Fig S1b) and had no relationship with the sedative-like effect of CCPA (Fig S1d).

Rats were subjected to 28 days of CUS exposure followed by SPT to screen out the depression-like ones. The depression-like rats were divided into four groups and were single intraperitoneally injected with CCPA 0 mg/kg, 0.002 mg/kg, 0.02 mg/kg, 0.2 mg/kg respectively, the control group were injected with vehicle 2 ml/kg intraperitoneally. The depression-like behaviors were assessed at 6 h, 12 h, 24 h, 36 h, 48 h, and 72 h after CCPA injection (Fig. S2a). Compared with the control group, CUS with CCPA 0 mg/kg group showed significant decrease in sucrose preference values (Fig. S2b) and increase in the immobility time in the FST (Fig. S2c), showing depression-like behaviors. After administration of CCPA (0.2 mg/kg), it rescued the CUS-induced decrease of sucrose preference value at 12 h, 24 h and 36 h, respectively. Meanwhile, CCPA (0.2 mg/kg) also rescued the CUS-induced increase of immobility time at 12 h, 24 h, 36 h and 48 h, respectively. Therefore, the depression-like behaviors induced by CUS were rapidly ameliorated at 12 h after injection of CCPA (0.2 mg/kg) and this effect persisted at least 36 hours. We conducted locomotor activity (LA) test to explore the effect of CCPA on LA in CUS rats. Results showed that the rapid antidepressant-like effects of CCPA (0.2 mg/kg) were independent of its effect on LA (Fig. S3b).

In addition, we also tested the anxiolytic-like effects of CCPA. In the OFT and the EPM test, CCPA rescued the significant decrease of the time to stay at central zone and to stay at open arms induced by CUS at from 7 h to 48 h after intraperitoneal injection (Figure S3c, d). And this anxiolytic-like effect of CCPA was also independent of its effect on LA (Figure S3b). See Supplementary Information for details.

To explore the mechanism underlying the depression-like behaviors induced by CUS, we detected the A_1_R signaling pathway and its downstream MAPK pathway [12]. Rats were randomly divided into a CUS group and a control group, and they were subjected to 28 days of CUS exposure or handling respectively, followed by sucrose preference test to screen out the successfully modeled rats. Then each group was divided randomly into two subgroups to receive injection of vehicle or CCPA at 9:00 am (ZT 13), and 11 hours later (ZT 0) the brain tissues were collected. Western Blot showed that in the hippocampal CA1 brain region, *p*-CREB and PER1 levels were decreased, which could be rescued by CCPA (Fig. 1b). Comparatively, in the mPFC region, *p*-CREB level was also decreased, however, this phenomenon could not be rescued by CCPA (Fig. 1c). Meanwhile, PER1 expression was not significantly altered in mPFC in the CUS group (Fig. 1c). In addition, we also did not find any effect of CUS or CCPA on the expression of A_1_R in these two brain regions (Fig. 1d and 1e).

**Fig. 1.**
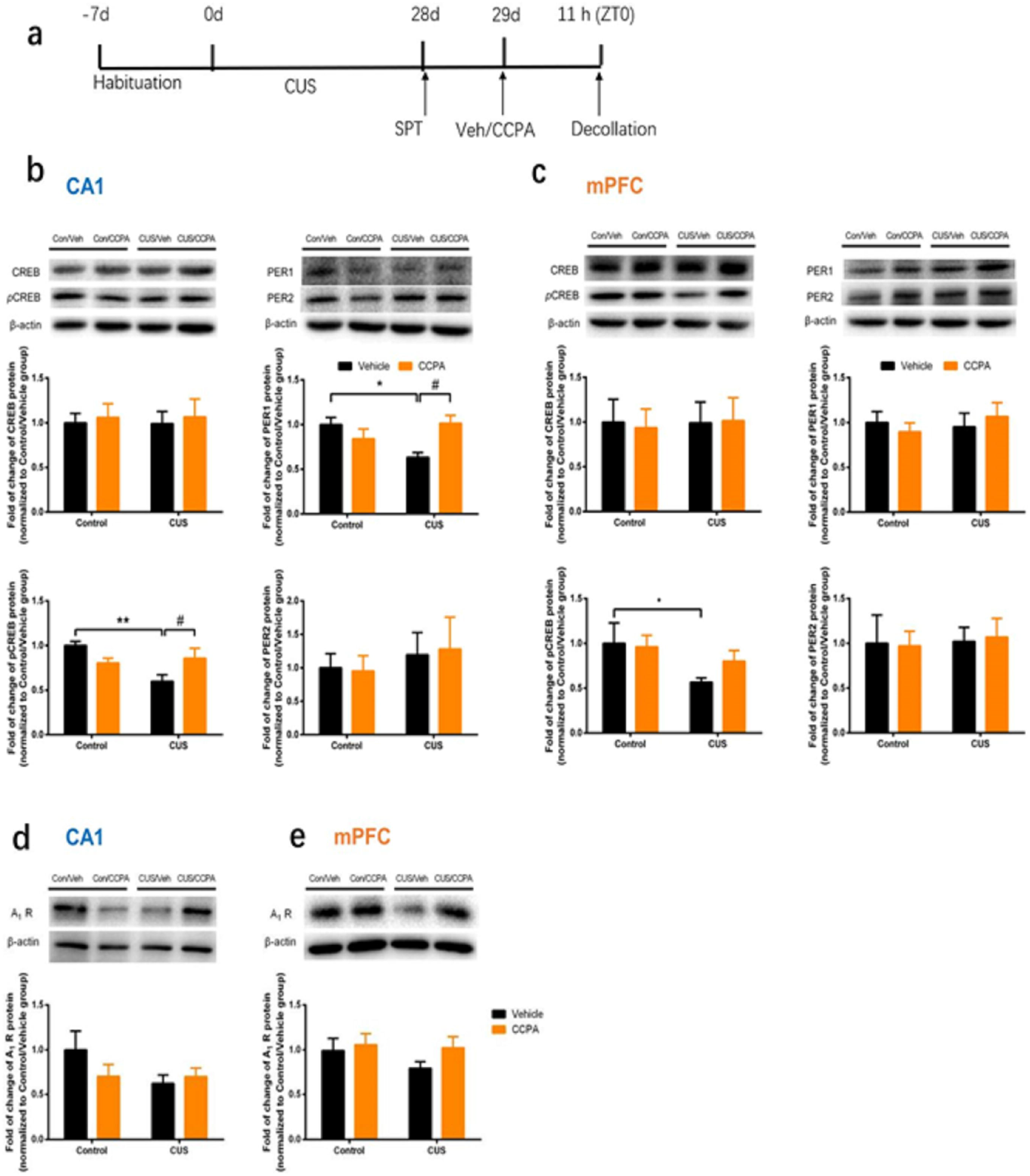
CUS induced decrease of *p*-CREB and PER1 in hippocampal CA1 could be rescued by CCPA. **a.** Timeline of the experiment procedure. **b.** Representative western blots and quantification of fold changes in CREB, *p*-CREB, PER1 and PER2 in the hippocampal CA1 region 11 h (ZT 0) after a single CCPA intraperitoneal injection in rats. *n* = 6 per group, two-way ANOVA, *p*-CREB: *F* stress _[1, 20]_ = 5.271, *P* < 0.05; *F* treatment _[1, 20]_ = 0.1828, *P* > 0.05; *F* stress × treatment _[1, 20]_ = 8.989, *P* < 0.01 and followed by *post hoc* Bonferroni’s test, * *P* < 0.05, ** *P* < 0.01, vs. control + veh group; ^#^ *P* < 0.05 vs. CUS + veh group. PER1: *F* stress _[1, 20]_ = 1.282, *P* > 0.05; *F* treatment _[1, 20]_ = 1.689, *P* > 0.05; *F* stress × treatment _[1, 20]_ = 10.02, *P* < 0.01 and followed by *post hoc* Bonferroni’s test, * *P* < 0.05, vs. control + veh group, ^#^ *P* < 0.05 vs. CUS + veh group. **c.** Representative western blots and quantification of fold changes in CREB, *p*-CREB, PER1 and PER2 in the mPFC 11 h (ZT 0) after a single CCPA intraperitoneal injection in rats. *n* = 6 per group, two-way ANOVA, *p*-CREB: *F* stress _[1, 20]_ = 4.359, *P* < 0.05; *F* treatment _[1, 20]_ = 0.5166, *P* > 0.05; *F* stress × treatment _[1, 20]_ = 0.9790, *P* > 0.05. **d.** Expression levels of adenosine A1 receptors in rat CA1. *n* = 6 per group, two-way ANOVA. **e.** Expression levels of adenosine A_1_ receptors in rat mPFC. *n* = 6 per group, two-way ANOVA. Data were presented as mean±SEM.

These results suggested that the *p*-CREB-PER1 signaling pathway in hippocampal CA1 region might be involved in the pathogeneses of depression.

### Knockdown of *Per1* in hippocampal CA1 induced depression-like behaviors

To further investigate whether the clock gene *Per1* is involved in the pathogenesis of depression, we infused adeno-associated virus (AAV)-mediated short hairpin RNA into hippocampal CA1 region to knockdown *Per1* expression in rats. As expected, three weeks later, the microinjection of AAV-shPer1 decreased the expression of PER1 and sucrose preference values significantly, while increased the immobility time in FST significantly, compared with the microinjection of AAV-Scramble group (Fig. 2b-e). The depression-like behaviors were relieved 12 h after injection of CCPA and this effect lasted until 36 h after injection (Fig. 2g, h). Western blotting showed that CCPA (0.2 mg/kg, i.p.) rescued the down-regulation of PER1 level caused by AAV-sh*Per1* (Fig. 2c).

**Fig. 2.**
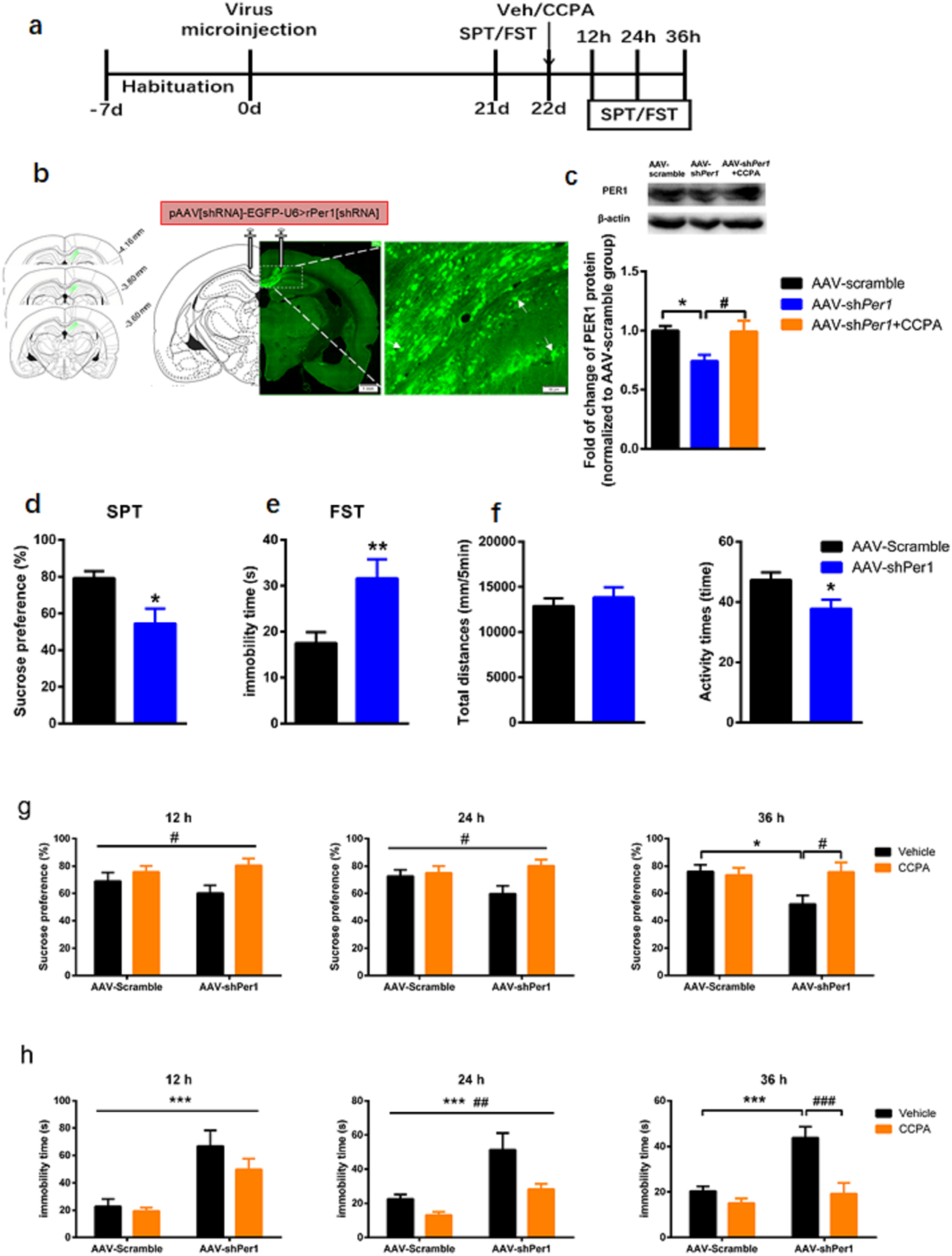
Knockdown the expression of *PER1* in hippocampal CA1 induced depression-like behaviors, which could be rescued by CCPA. **a.** Timeline of the experiment procedure. **b.** Immunofluorescence showed the injection sites and coronal section of the hippocampal CA1 region. Scale bars = 500 um (low-magnification images) on the up-right panel, Scale bars = 20 um (high-magnification images) on the down-right panel. **c.** PER1 expression in the CA1 in rats that were microinfused with AAV-Scramble or AAV-sh*Per1*, and AAV-sh*Per1* group injected with CCPA (i.p.), quantified by western blot (*n* = 5 per group). unpaired *t* - test, *t* = 2.893, * *P* < 0.05; Mann Whitney test, u = 2.000, ^#^ *P* < 0.05. **d.** Sucrose preference test (SPT), *n* = 10 per group, Mann Whitney test, u = 23, * *P* < 0.05 vs. AAV-Scramble group. **e.** Forced swimming test (FST), *n* = 10 per group, unpaired *t* - test, *t* = 2.986, ** *P* < 0.01 vs. AAV-Scramble group. **f.** Locomotor activity tests showed that AAV-sh*Per1* group decreased significantly in activity times (*n* = 10 per group, unpaired *t* test, *t* = 2.419, * *P* < 0.05), not in total distances. **g.** SPT at 12-36 h after a single intraperitoneal injection of vehicle or CCPA. *n* = 9 - 10 per group, two-way ANOVA, 12 h: Vehicle vs. CCPA, *F* treatment _[1, 35]_ = 6.255, ^#^ *P* < 0.05; *F* sh*Per1* _[1, 35]_ = 0.1346, *P* > 0.05; *F* treatment × sh*Per1* _[1, 35]_= 1.514, *P* > 0.05; 24 h: *F* treatment _[1, 36]_= 5.265, ^#^ *P* < 0.05; *F* sh*Per1* _[1, 36]_= 0.6054, *P* > 0.05; *F* treatment × sh*Per1* _[1, 36]_= 3.317, *P* > 0.05; 36 h: *F* treatment _[1, 36]_ = 3.175, *P* > 0.05; *F* sh*Per1* _[1, 36]_ = 3.390, *P* > 0.05; *F* treatment × sh*Per1* _[1, 36]_ = 4.830, *P* < 0.05 and followed by *post hoc* Bonferroni’s test, * *P* < 0.05, vs. AAV-Scramble + Vehicle group, ^#^ *P* < 0.05, vs. AAV-sh*Per1* + CCPA group. **h.** FST at 12 - 36 h after a single intraperitoneal injection of vehicle or CCPA. *n* = 10 per group, two-way ANOVA, 12 h: *F* treatment _[1, 36]_= 1.877, *P* > 0.05; *F* sh*Per1* _[1, 36]_= 24.98, *P* < 0.0001; *F* treatment × sh*Per1* _[1, 36]_= 0.856, *P* > 0.05; 24 h: *F* treatment _[1, 36]_= 9.236, ^##^*P* < 0.01; *F* sh*Per1* _[1, 36]_= 17.17, \*\*\**P* < 0.001; *F* treatment × sh*Per1* _[1, 36]_ = 1.673, *P* > 0.05; 36 h: *F* treatment _[1, 36]_ = 16.20, *P* < 0.001; *F* sh*Per1* _[1, 36]_ = 13.64, *P* < 0.001; *F* treatment × sh*Per1* _[1, 36]_ = 6.737, *P* < 0.05 and followed by *post hoc* Bonferroni’s test, \*\*\**P* < 0.001 vs. AAV-Scramble + Vehicle group, ^###^*P* < 0.001 vs. AAV-sh*Per1* + CCPA group. Data were presented as Mean ± SEM.

These results indicated that *Per1* in hippocampal CA1 might play a key role in the pathogenesis of depression and mediate the rapid antidepressant-like effect of CCPA.

### Knockdown of *Per2* in hippocampal CA1 showed potential antidepressant-like effect

Both clock genes *Per1* and *Per2* have CRE sequences (TGACGTCA) in their promotor regions [16], which are the binding sites of *p*-CREB. We further explored whether the clock gene *Per2* was involved in the pathogenesis of depression. In the same way, AAV-scramble or AAV-sh*Per2* was microinjected into hippocampal CA1 in rats. Three weeks later, the expression of PER2 in hippocampal CA1 region significantly decreased in AAV-sh*Per2* group, compared with that in the AAV-Scramble group (Fig. 3b and 3c). Contrary to the results of AAV-sh*Per1*, the immobility time of the AAV-sh*Per2* group was significantly decreased in FST compared with the AAV-Scramble group, indicating that the down-regulation of PER2 resulted in a potential antidepressant-like effect (Fig. 3e).

**Fig. 3.**
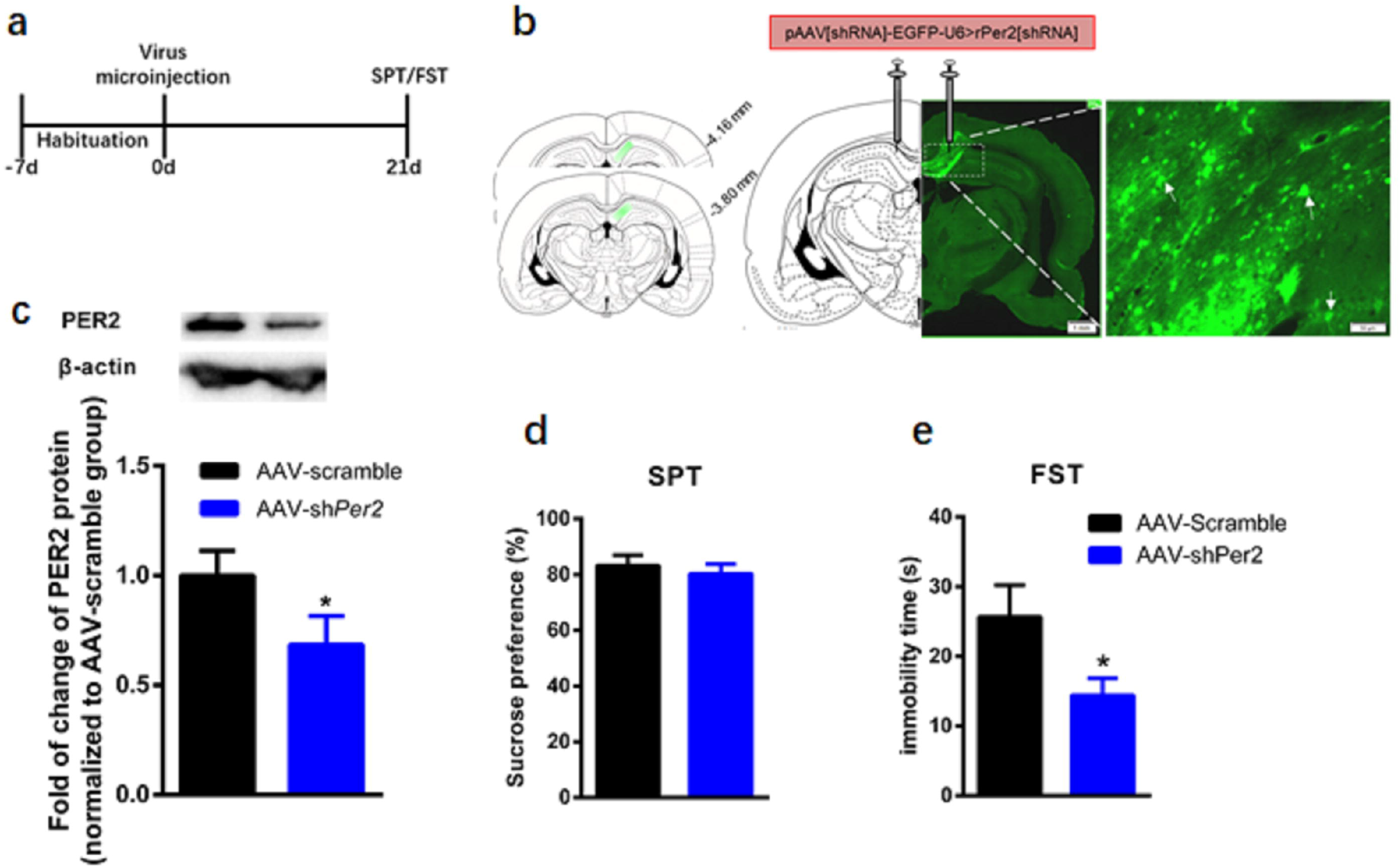
Knockdown the expression of *PER2* in hippocampal CA1 exhibited antidepressant-like effect potentially. **a.** Timeline of the experiment procedure. **b.** Immunofluorescence showed the injection sites and coronal section of the hippocampal CA1 region. Scale bars = 500 um (low-magnification images) on the up-right panel, Scale bars = 20 um (high-magnification images) on the down-right panel. **c.** PER2 expression in the CA1 in rats that were microinfused with AAV-Scramble or AAV-sh*Per2*, quantified by western blot, *n* = 4 per group, unpaired *t*-test, *t* = 2.494, * *P* < 0.05, vs. AAV-Scramble group. **d.** Sucrose preference test (SPT), *n* = 10 per group, unpaired *t*-test, *t* = 0.2556, *P* > 0.05. **e.** Forced swimming test (FST). *n* = 10 per group, unpaired *t* - test, *t* = 2.374, * *P* < 0.05. Data were presented as mean±SEM.

These results suggested that both *Per1* and *Per2* in hippocampal CA1 play key roles in the pathogenesis of depression. In hippocampal CA1, *Per1* may be a protective factor and *Per2* may be a risk factor for the occurrences of the depression-like behaviors in rats.

### Knockout the CRE sequence in the promoter region of either *Per1* or *Per2* led to depression-like behaviors

As far as we know, CRE sequences exist in the promoter regions of hundreds of genes [13, 14, 16]. The combination of *p*-CREB with the CRE sequence in the promoter regions of *Per1* and *Per2* may be an important path in the pathogenesis of depression and it also may be a key target of CCPA. We used rats with point mutations (Del_TGACGTCA) on the promoter region of *Per1* or *Per2* locus by CRISPR/Cas9-mediated genomic engineering to further confirm our hypothesis. The schematic diagram of the targeting strategy of rat *Per1 or Per2* (Del_TGACGTCA) knock-in project was shown in Fig. S 4a and S 4b.

Sanger sequencing results showed that the CRE sequence (TGACGTCA) at the rat *Per1* or *Per2* locus was deleted (took No.60 as an example), compared to the wild-type sequence (Fig. S4c and S4d). After the homozygotes and the wild-type grew for at least 8 weeks, the following experiments can be performed after verification. Compared to the wild-type group, the sucrose preference values of the homozygotes of both the *Per1* CRE sequence deletion group and the *Per2* CRE sequence deletion group were significantly decreased (Fig. 4b), while the immobility time of the homozygotes of *Per2* CRE sequence deletion group was significantly increased in the FST (Fig. 4c). As expected, CCPA couldn’t rescue these depression-like behaviors (Fig. 4d and 4e). We performed western blot to further explore the effect of CRE sequence on the expression of PER1 and PER2 in hippocampal CA1 region. The results showed that the PER2 level in the hippocampal CA1 region was significantly increased in the homozygous of *Per1* gene CRE knockout group, compared to that in the wild-type group (Fig. 4f).

**Fig. 4.**
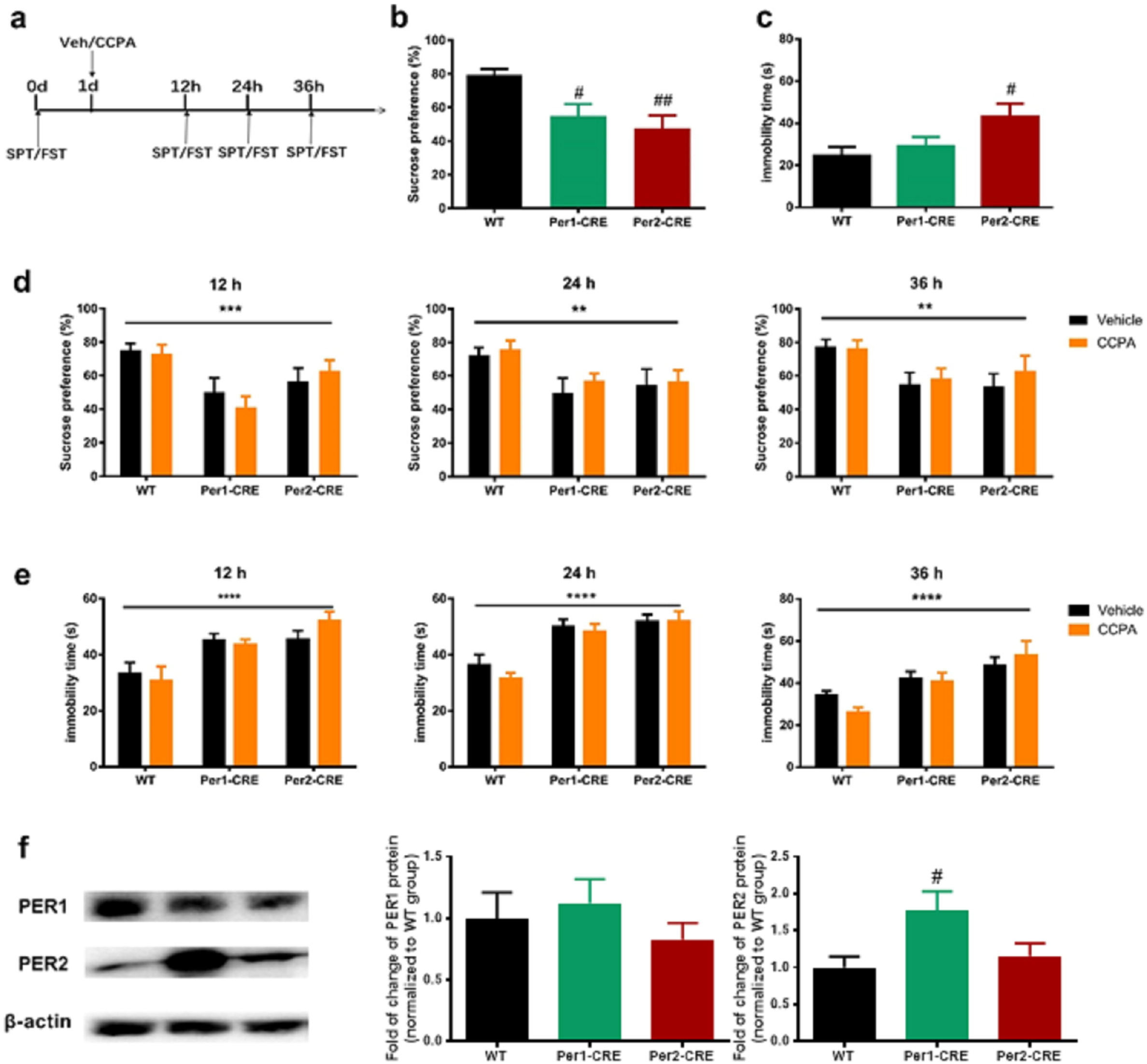
CRE sequence in the promotor region of *Per1* or *Per2* deletion resulted in depression-like behaviors, which could not be rescued by CCPA. **a.** Timeline of the experiment procedure. **b.** Sucrose preference test (SPT) before a single intraperitoneal injection of vehicle or CCPA. *n* = 10 per group, one-way ANOVA, *F* _2, 27_ = 6.917, *P* < 0.01, followed by *post hoc* Bonferroni’s test, ^#^*p* < 0.05, ^##^ *p* < 0.01, vs. wild type (WT) group. **c.** Forced swimming test (FST) before a single intraperitoneal injection of vehicle or CCPA. *n* = 10 per group, one - way ANOVA, *F* _2, 27_ = 3.804, *P* < 0.05, followed by *post hoc* Bonferroni’s test, ^#^ *p* < 0.05, vs. WT group. **d.** SPT after 12 - 36 h a single intraperitoneal injection of vehicle or CCPA. *n* = 9 - 10 per group, two - way ANOVA, 12 h: *F* CRE _[2, 53]_= 9.044, *** *P* < 0.001; *F* treatment _[1, 53]_= 0.063, *P* > 0.05; *F* CRE × treatment _[2, 53]_= 0.6865, *P* > 0.05; 24 h: *F*CRE _[2, 54]_=5.508, ** *P* < 0.01; *F* treatment _[1,54]_= 0.5421, *P* > 0.05; *F* CRE × treatment _[2, 54]_= 0.06642, *P* > 0.05; 36 h: *F* CRE _[2, 54]_=5.773, ** *P* < 0.01; *F* treatment _[1, 54]_= 0.5095, *P* > 0.05; *F* CRE × treatment _[2, 54]_= 0.3031, *P* > 0.05. **e.** FST after 12 - 36 h a single intraperitoneal injection of vehicle or CCPA. *n* = 10 per group, two - way ANOVA, 12 h: *F* CRE _[2, 54]_= 12.83, **** *P* < 0.001; *F* treatment _[1, 54]_= 0.098, *P* > 0.05; *F* CRE × treatment _[2, 54]_ = 1.093, *P* > 0.05; 24 h: *F* treatment _[2, 54]_ =22.33, **** *P* < 0.0001; *F* treatment _[1, 54]_ = 0.8133, *P* > 0.05; *F* CRE × treatment _[2, 54]_= 0.3558, *P* > 0.05; 36 h: *F* CRE _[2, 54]_ = 12.75, **** *P* < 0.0001; *F* treatment _[1, 54]_ = 0.1781, *P* > 0.05; *F* CRE × treatment _[2, 54]_ = 1.301, *P* > 0.05. **f.** Western blot showed the PER1 and PER2 expression levels in hippocampal CA1 region in WT, *Per1* CRE-knockout and *Per2* CRE-knockout group. *n* = 5 per group, one - way ANOVA, PER1: *F* _2, 12_ = 0.6447, *P* > 0.05; PER2: *F*_2, 12_ = 4.979, *P* < 0.05 followed by *post hoc* Bonferroni’s test, ^#^*P* < 0.05, vs. WT group. Data were presented as mean ± SEM.

These results indicated that the CRE sequence in the promoter region of *Per1* and *Per2* is essential in the pathogenesis of depression and is indispensable in the antidepressant-like effect of CCPA.

In summary, these results demonstrated that the A_1_R-*p*-CREB-*Per1/2* pathway in the CA1 region in rats may be involved in the pathophysiology of depression and mediate the rapid antidepressant-like effect of A_1_R agonist CCPA, as shown in Fig. 5.

**Fig.5.**
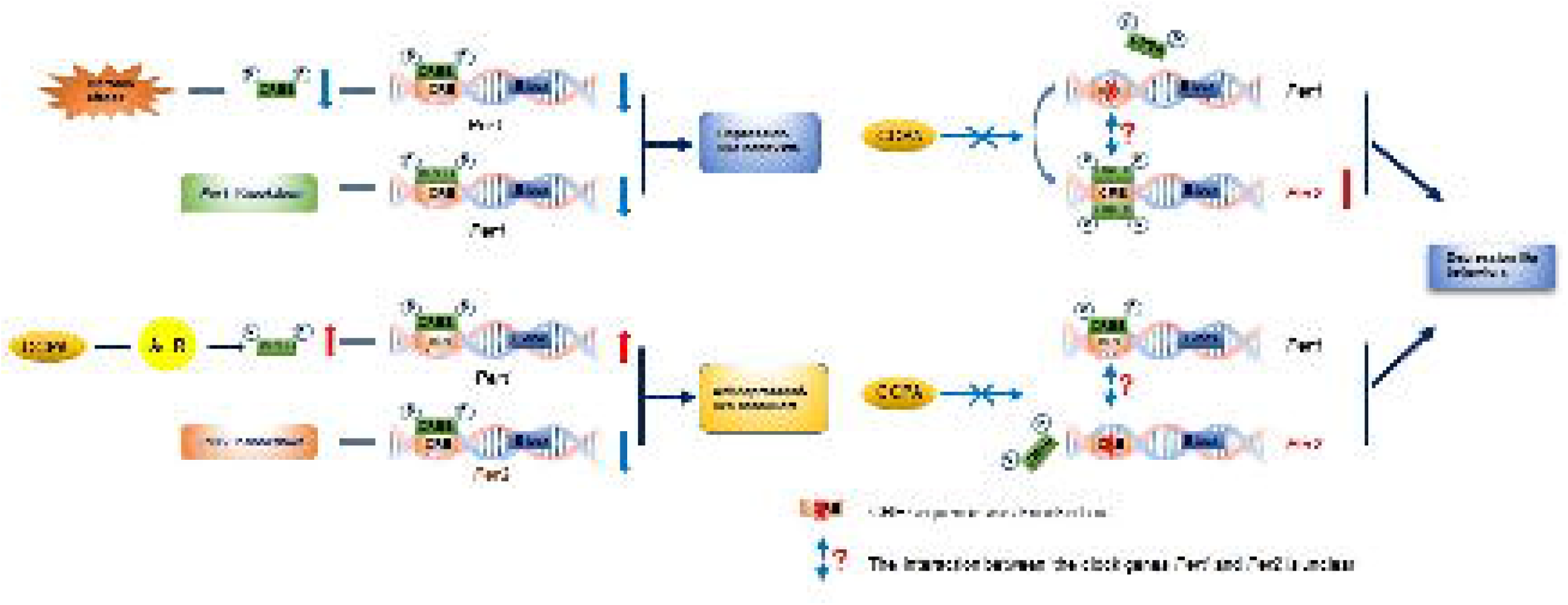
Schematic diagram that illustrates the involvement of clock genes *Per1* and *Per2* in the hippocampal CA1 in depression-like behaviors and antidepressant response. Left: Chronic stress decreased *p*-CREB and PER1 levels in the hippocampal CA1 and induced depression - like behaviors. Simultaneously, the knockdown of PER1 levels in CA1 also induced depression-like behaviors. Adenosine A1 receptor agonist CCPA rescued the chronic stress-induced decrease in *p*-CREB and PER1 expression in CA1 and produced rapid antidepressant - like effect. *Per2* knockdown resulted in potentially antidepressant - like effect. Right: Both homozygotes of SD-*Per1* (c.-3353_-3345delTGACGTCA) Smoc and SD-*Per2* (c.-10368_-10360delTGACGTCA) Smoc showed depression - like behaviors, which could not be rescued by CCPA. Furthermore, the PER2 gene expression levels in CA1 in homozygotes of SD-*Per1* (c.-3353_-3345delTGACGTCA) Smoc significantly increased. A1R, adenosine A1 receptor.

## Discussion

In this study, we demonstrated that CUS induced the depression-like behaviors and the decrease of the expression of *p*-CREB and PER1 in CA1 in rats, which could be rapidly rescued by CCPA. *Per1* knockdown in the hippocampal CA1 region induced depression-like behaviors, which could be also rapidly rescued by CCPA. However, *Per2* knockdown in the hippocampal CA1 region led to potential antidepressant-like behaviors. Then we deleted the CRE sequence in the promotor regions in *Per1* or *Per2*, the homozygous rats showed depression-like behaviors, but CCPA could not rescue these depression-like behaviors.

Behavioral studies showed that CCPA exhibited an antidepressant-like effect from 12 h to at least 36 h after injection, consistent with the previous study [8]. Our results that CCPA had a sedative-like effect in the first few hours after injection were also consistent with the previous study [27]. The antidepressant-like effect of CCPA was independent of its effect on locomotor activity.

In line with previous study [28], the expression of *p*-CREB and PER1 in hippocampal CA1 region was down-regulated by CUS, which could be rescued by CCPA. Nevertheless, the expression of PER2 in CA1 region was not altered by CUS. Previous study reported that the level of *Per2* mRNA was significantly increased in CA1 region in CUS rats compared to the control group [28]. This difference may be due to our research at the protein level rather than at the mRNA level. It is well known that there are many alterations of circadian genes in the process from RNA to protein expression, including alternative splicing, RNA degeneration, protein phosphorylation, ubiquitination and other post-translational modifications [29].

Knockdown of *Per1* in CA1 region led to depression-like behaviors, which could also be rescued by CCPA. This phenomenon might be related to the reversal of PER1 expression by CCPA. In contrast, knockdown of *Per2* in CA1 region showed potential antidepressant-like effect. These results indicated that *Per1* in CA1 region might be a protective factor and *Per2* in CA1 region might be a risk factor for depression-like behavior in rats. Under normal circumstances, the two factors maintain a balance in the CA1 region, thus ensuring the normal behaviors of rats. When exposed to CUS, the expression of PER1, a protective factor in the CA1 region, was down-regulated and PER2 protein level remained unchanged. The balance between the two factors was disturbed, the rats would exhibit depression-like behaviors. Similarly, when the expression of *Per2* in CA1 region was down-regulated, the risk factor was reduced and the rats exhibited antidepressant - like behaviors.

In order to further explore the targets of *p*-CREB acting through the clock gene *Per*, we constructed model rats, in which we used CRISPR-Cas9 technique to knock out the CRE sequence on the promoter of *Per1* or *Per2* [16]. We found that homozygotes with knockout of CRE sequence on the promoter region of *Per1* or *Per2* gene showed depression - like behaviors. This result indicates that the CRE sequence on the promoter region in *Per1/2* may be a key point in the pathogenesis of depression-like behaviors in rats.

Besides, the deletion of the CRE sequence didn’t affect the expression of both genes, although the deletion of the CRE sequence in the *Per1* gene significantly increased the level of Per2 protein. In line with a previous study, we found that the PER2 level in *Per2* homozygous rats was not affected by the CRE deletion [30]. However, the PER2 level in *Per1* homozygous rats was significantly increased due to the CRE deletion. There was no significant change in the PER1 level in either *Per1* or *Per2* homozygous rats. The PER1 level was in line with the results before light pulse in the previous study [30]. Our result of the PER2 level in *Per1* homozygous rats may be due to the different brain region and the different purpose we studied.

As expected, CCPA could not rescue the depression-like behaviors of these CRE deleted homozygous rats, indicating that the CRE sequences on the promoter regions in *Per1* and *Per2* genes play indispensable roles underlying the mechanism of antidepressant-like effect of CCPA.

According to the western blot results, the deletion of CRE sequence in *Per1* and *Per2* genes had no effect on the protein level of PER1. Moreover, the deletion of the CRE sequence in *Per2* also didn’t change the PER2 protein level. One reason may be that there are many promoters in the *Per* gene, such as E-box, glucocorticoid-responsive elements (GRE). Only knockout of the CRE sequence on the promoter region cannot influence the expression of the gene itself. Another reason may be that the CRE sequence on the *Per2* gene is not involved in the PER1 expression process. However, the expression of PER2 was significantly increased in *Per1* homozygous rats with CRE sequence knockout on the promoter region. This might indicate that the CRE sequence on the promoter region in *Per1* gene is important for the PER2 expression process and might have an inhibitory or complementary effect. Once the CRE sequence on the promoter region in the *Per1* gene is deleted, the *Per2* gene may increase its expression level through its own CRE. Therefore, we deduced the following possibilities. First, clock gene *Per1* and *Per2* might compete in binding with *p*-CREB via their CRE sequence. Second, clock gene *Per1* and *Per2* may be complementary in the pathogenesis of depression, and the CRE sequence may play a critical role in their relationship. Third, the expression of PER2 may be regulated by *Per1* through the CER sequence on the promoter region, while the expression of PER1 may not be affected by the CRE sequence on the *Per2* gene. There may be other molecules involved in this process. All in all, further investigations are needed to explore the detailed relationship between them.

As for limitations, first, our results mainly studied the role of clock gene *Per1/2* in the pathogenesis of depression and the mechanism underlying the antidepressant-like effect of CCPA from the perspective of non-rhythmic system. The part of roles of the CREs on *Per1* or/and *Per2* in the circadian system have been reported in a recent research [30]. But their interaction in the circadian rhythm remains to be elucidated in the future. Second, our research only used CCPA as an antidepressant, and the key role of the CRE sequence on the promoter region in the *Per* gene cannot be extended to other antidepressant strategies. We predict here that almost all antidepressant strategies, especially rapid antidepressant therapies, may act through the combination of *p*-CREB with the CRE sequence on the promoter region in the *Per* gene. This hypothesis needs further research in the future.

## Conclusions

Our research revealed that the clock genes *Per1* and *Per2* in the hippocampal CA1 region in rats are involved in the pathogenesis of depression-like behavior and mediate the rapid antidepressant-like effect of CCPA. The CRE sequence in the promoter region of clock genes *Per1/2* is critical and may be a target for the development of novel rapid antidepressants in the future.

## Supporting information

https://pan.baidu.com/s/1i3wnF7qqHPeEUDnmKWRXIw Extracted Number: jtqt

## Acknowledgments

We thank Professor George Fu Gao and Lin Lu for their supervision and guidance. They gave us a lot of advice, supplies and help in the design and performance throughout the whole research. This study was supported by the National Natural Science Foundation of China (81871071, 81171251, 81901352, 82001404 and 32071058), the Natural Science Foundation of Beijing Municipality, China (7162101) and Peking University Medicine Fund of Fostering Young Scholars’ Scientific & Technological Innovation and the Fundamental Research Funds for the Central Universities (No. BMU2020PYB013).

