## Supplementary material for "Clock genes *Period 1* and *Period 2* in the hippocampal CA1 mediate depression-like behaviors and rapid antidepressant response": https://pan.baidu.com/s/1i3wnF7qqHPeEUDnmKWRXIw Extracted Number: jtqt

Running Title: Clock gene *Per* mediates depression-like behaviors

Abstract: 239 words

Introduction:	467 words

Discussion: 1095 words

Total text: 3316 words

Figures: 5

*Corresponding author:

Su-Xia Li, MD., PhD

National Institute on Drug Dependence, Peking University

38, Xue Yuan Road, Haidian District, Beijing 100191, China

**Acknowledgments**

We thank Professor George Fu Gao and Lin Lu for their supervision and guidance. They gave us a lot of advice, supplies and help in the design and performance throughout the whole research. This study was supported by the National Natural Science Foundation of China (81871071, 81171251, 81901352, 82001404 and 32071058), the Natural Science Foundation of Beijing Municipality, China (7162101) and Peking University Medicine Fund of Fostering Young Scholars’ Scientific & Technological Innovation and the Fundamental Research Funds for the Central Universities (No. BMU2020PYB013).

**Declaration of interests**

The authors declare no conflict of interest.

**Materials and Methods**

***Sucrose preference test and forced swim test***

The procedures were based on former studies (Banasr and Duman, 2008; Shi et al., 2012). After adaptation for 48 h, the rats were deprived of water and food for 4 h and then subjected to the sucrose preference test, in which they were housed in individual cages for 1 h and had free access to two bottles that contained 1% sucrose or tap water. The forced swim test (FST) was undergone in a plastic cylinder (25 cm diameter and 65 cm height) that was filled with water (45 cm depth) at a temperature of 23–25 °C. Immobility was defined as minimal movement of both the four limbs and tail of the rats.

***Chronic unpredictable stress (CUS) protocol***

CUS was undertaken with variable sequences of different mild stressors, with two stressors per day for 28 days, which was revised from previous studies (Banasr and Duman, 2008; Li et al., 2017). Rats were exposed to chronic unpredictable stress with two stressors per day for 28 days. All the stressors included restraint (immobilization, 1 h in the dark period from ZT 12 to ZT 24), forced swim (swim in 10 °C water, 5 min in the light period from ZT 0 to ZT 12), cage rotation (rat cages were placed in a shaker, 120 rpm/min, 1 h during the dark phase from ZT 12 to ZT 24), cold (rats were put into an ice box with ventilation at 4 °C, 1 h in the dark period), damp bedding (rats stayed with wet bedding, overnight), cage titling (the cage was placed at 45° for 24 h), crowding (12 rats in a cage at size of 41 × 26 × 21 cm, overnight), strobe light (rats were put into a box with white flash light of 10-100 lux at 2 Hz for 6 h from ZT 0 to ZT 6 ), empty cages (removal of bedding from the cages for 24 h) , noise (rats were kept in the room with 120 dB white noises for 1 h in the dark phase), water deprivation (24 h), and food deprivation (24 h). Two stressors were given to the rats in a variable sequence every day. Considering the effect of light-dark reversal (24 h) on diurnal rhythm, we didn’t use this stressor in our protocols. In parallel, control rats were maintained under the same environment without any stressors, with the exception of daily handling.

***Locomotor activity***

Locomotor activity (LA) was measured with an automated video-tracking system (DigBehv-LM4; Shanghai Jiliang Software Technology), which had been used in our previous studies (Sun et al., 2013). A monochrome video camera was mounted on top of each chamber. All of the chambers were connected to a computer. The video files (stored on the computer) were analyzed using DigBehv analysis software. Locomotor activity was expressed as the total distance traveled during the 5 min test.

***Open field test***

The open field apparatus was a 75 cm wide × 75 cm long × 45 cm high arena with brown plywood walls and a wooden floor. The floor was divided into 25 equal 15 × 15 cm squares by black lines. The rats were placed in the box to explore the novel environment for 5 min and then returned to their home cage. To familiarize the rats with the arena, they were placed in the arena for 30 min 1 d before the experiment (Ballarini et al., 2009; Moncada and Viola, 2006).

***Elevated plus maze***

The elevated plus maze test was based on our previous studies (Suo et al., 2013). The entire test was conducted under dim light conditions. Each rat was first placed in the central zone of the elevated plus maze and allowed to freely explore it for 5 min. The number of entries into and time spent (in seconds) on the open arms were recorded by two independent observers who were blind to the animal groups.

***Chronic unpredictable stress procedure***

The protocol was adapted from previous studies (Jiang et al., 2013; Katz et al., 1981). Rats were exposed to chronic unpredictable stress with two stressors per day for 28 days. All the stressors included restraint (immobilization, 1 h in the dark period from ZT 12 to ZT 24), forced swim (swim in 10 °C water, 5 min in the light period from ZT 0 to ZT 12), cage rotation (rat cages were placed in a shaker, 120 rpm/min, 1 h during the dark phase from ZT 12 to ZT 24), cold (rats were put into an ice box with ventilation at 4 °C, 1 h in the dark period), damp bedding (rats stayed with wet bedding, overnight), cage titling (the cage was placed at 45° for 24 h), crowding (12 rats in a cage at size of 41 × 26 × 21 cm, overnight), strobe light (rats were put into a box with white flash light of 10-100 lux at 2 Hz for 6 h from ZT 0 to ZT 6 ), empty cages (removal of bedding from the cages for 24 h) , noise (rats were kept in the room with 120 dB white noises for 1 h in the dark phase), water deprivation (24 h), and food deprivation (24 h). Two stressors were given to the rats in a variable sequence every day. Considering the effect of light - dark reversal (24 h) on diurnal rhythm, we didn’t use this stressor in our protocols. In parallel, control rats were maintained under the same environment without any stressors, with the exception of daily handling.

**Results**

**The potential antidepressant-like effects of CCPA**

We used forced swimming to test whether CCPA has a potential antidepressant-like effects. Rats were randomly divided into four groups and intraperitoneally injected with vehicle or CCPA (0.002, 0.02 and 0.2 mg/kg), respectively. FSTs were performed at 6 h, 12 h, 24 h, 36 h, 48 h, and 72 h after injection (Figure S1a). The immobility time decreased significantly in the CCPA (0.02 and 0.2 mg/kg) group compared with control group at 24 h and persisted at least to 36 h after injection (Figure S1b). To exclude the effect of CCPA on locomotor activity, we performed LA tests with rats injected with vehicle or CCPA (0.2 mg/kg). The results showed that CCPA (0.2 mg/kg) had a sedative-like effect compared with vehicle at 5 h and disappeared at 12 h after injection (Figure S1d). These results suggested that the decreased immobility time in the FST persisted from 24 h to 36 h had no relationship with the sedative-like effect of CCPA.

**CUS induced depression- and anxiety-like behaviors could be rapidly rescued by CCPA**

To further confirm the antidepressant-like effects of CCPA, rats subjected to 28 days of CUS exposure or handling, followed by SPT to screen out the depression-like ones. The depression-like rats were divided into four groups and were intraperitoneally injected with CCPA 0 mg/kg, 0.002 mg/kg, 0.02 mg/kg, 0.2 mg/kg respectively, the control group were injected with vehicle 2 ml/kg intraperitoneally. The depression-like behaviors were assessed at 6 h, 12 h, 24 h, 36 h, 48 h, and 72 h after injection (Fig.S2a). Compared with control group, CUS + CCPA 0 mg/kg group showed significant decrease in sucrose preference value (Fig. S2b) and increase in immobility time in the FST (Fig. S2c), showing depression-like behaviors. CCPA (0.2 mg/kg) rescued the CUS induced decrease of sucrose preference value at 12 h, 24 h and 36 h, respectively. Meanwhile, CCPA (0.2 mg/kg) also rescued the CUS induced increase of immobility time at 12 h, 24 h, 36 h and 48 h, respectively. Therefore, the depression - like behaviors induced by CUS were rapidly ameliorated at 12 h after injection of CCPA (0.2 mg/kg) and persisted at least 36 h.

To explore the effect of CCPA on LA in CUS rats, we conducted LA test with rats injected with vehicle or CCPA (0.2 mg/kg). Results showed that CCPA (0.02 mg/kg) enhanced the LA compared with vehicle at 5 h and disappeared at 12 h after injection (Figure S3b). This suggests that the rapid antidepressant - like effects of CCPA (0.2 mg/kg) was independent of its effect on LA.

In addition, we also tested the anxiolytic-like effects of CCPA. In the OFT and the EPM test, rats spent the time to stay at central zone and to stay at open arms significantly decreased in CUS group compared to that in the control group, showing anxiety - like behaviors induced by CUS (Figure S3c, d). CCPA rescued these anxiety-like behaviors at from 7 h to 48 h after intraperitoneally injection (Figure S2 b, c), demonstrating anxiolytic-like effects. And this anxiolytic-like effect of CCPA was also independent of its effect on LA (Figure S3b).

**Figure Legends**

**Fig. S1 CCPA exhibited potential antidepressant-like effect.**

**a.** Timeline of the forced swim test (FST). **b.** FST at 6 h, 12 h, 24 h, 36 h, 48 h, and 72 h after a single vehicle (veh) or CCPA (0.002mg/kg, 0.02mg/kg and 0.2 mg/kg) intraperitoneal injection in rats. *n* = 9-10 per group; One-way ANOVA (24 h: *F* _3, 36_ = 19.49, *P* < 0.0001; 36 h: *F*_3, 32_ = 7.340, *P* = 0.0007) followed by *post hoc* Bonferroni’s test, **P* < 0.05, ** *P* < 0.01, *** *P* < 0.001, **** *P* <0.0001, vs. the veh group. **c.** Timeline of the locomotor activity tests (LA). **d.** LA (including total distances and activity times) at 5 h and 24 h after a single injection of veh or CCPA (0.2 mg/kg). *n* = 8 per group, unpaired *t* test, 5 h: total distances, *t* = 5.635, *****P* < 0.0001, vs. the veh group; activity times, *t* = 1.881, *P* > 0.05; 24 h: total distances, *t* = 0.092, *P* > 0.05; activity times, *t* = 0.405, *P* > 0.05. Data were presented as mean ± SEM.

**Fig. S2** **CUS induced depression-like behaviors could be rapidly rescued by** **CCPA.**

**a** Timeline of CUS exposure and behavioral tests. **b** Sucrose preference test (SPT). *n* = 8-10 per group. unpaired two - tailed *t* tests were performed to compare CUS + CCPA (0 mg/kg) group with Control + Vehicle (veh) group (*t* = 3.623, ** *P* < 0.01 at 6 h; *t* = 3.531, ** *P* < 0.01 at 12 h; *t* = 2.457, * *P* < 0.05 at 24 h; *t*= 3.533, *** P* < 0.01 at 36 h; *t* = 2.482, * *P* < 0.05 at 48 h; *t* = 2.361, ** P* < 0.05 at 72 h). One-way ANOVA followed by *post hoc* Bonferroni’s tests, 6 h: *F* _3,28_ = 0.2184, *P* > 0.05; 12 h: *F* _3, 28_ = 4.630, *P* < 0.01; 24 h: *F* _3, 28_ = 3.165, *P* < 0.05; 36 h: *F* _3, 28_ = 4.534, *P* < 0.05; 48 h: *F* _3, 28_ = 0.7626, *P* > 0.05; 72 h: *F* _3, 28_ = 0.9097, *P* > 0.05. ^#^ *P* < 0.05, ^##^ *P* < 0.01, vs. CUS + CCPA (0 mg/kg) group. **c** Force swimming test (FST). *n* = 8-10 per group. unpaired *t* tests were performed to compare CUS + CCPA (0 mg/kg) with Control + Veh group (*t* = 2.800, * *P* < 0.05 at 6 h; *t* = 2.209, * *P* < 0.05 at 12 h; *t* = 2.919, * *P* < 0.05 at 24 h; *t* = 5.340, *** *P* < 0.0001 at 36 h; *t* = 2.516, * *P* < 0.05 at 48 h; *t* = 2.466, * *P* < 0.05 at 72 h). One-way ANOVA followed by *post hoc* Bonferroni’s tests, 6 h: *F* _3, 36_ = 1.990, *P* > 0.05; 12 h: *F* _3, 36_ = 2.538, *P* = 0.072; 24 h: *F* _3, 28_ = 4.758, *P* < 0.01; 36 h: *F* _3, 28_ = 9.693, *P* = 0.0001; 48 h: *F* _3, 36_ = 22.09, *P* < 0.0001;72 h: *F* _3, 36_ = 1.719, *P* > 0.05. ^#^ *P* < 0.05, ^##^ *P* < 0.01, ^###^ *P* < 0.001, ^####^ *P*< 0.0001, vs. CUS + CCPA (0 mg/kg) group. Data were presented as Mean ± SEM.

**Fig. S3** **CUS induced anxiety-like behaviors could be rapidly rescued by** **CCPA.**

**a.** Timeline of the experiment procedure. **b.** Locomotor activity test (LA) at 5 h and 12 h after a single intraperitoneal injection of vehicle or CCPA. *n* = 8-10 per group, unpaired two-tailed *t* tests were performed to compare CUS + CCPA (0 mg/kg) group with Control + Vehicle group. * *P* < 0.05, ** *P* < 0.01, vs. control group. One-way ANOVA followed by *post hoc* Bonferroni’s test were performed to compare CUS groups that were given CCPA (0 mg/kg, 0.002 mg/kg, 0.02 mg/kg and 0.2 mg/kg) at 5 h and 12 h (*F*_3, 28_ = 7.041, *P* < 0.01 at 5 h in total distances; *F*_3, 28_ = 1.479, *P* > 0.05 at 5 h in activity times; *F*_3, 28_ = 0.8609, *P* > 0.05 at 12 h in total distances; *F*_3, 28_ = 0.6011, *P* > 0.05 at 12 h in activity times. ^###^ *P* < 0.001, vs. CUS + CCPA (0 mg/kg). **c.** Open field test (OFT) 7 h after a single intraperitoneal injection of vehicle or CCPA. *n* = 8-10 per group, unpaired two-tailed *t* tests were performed to compare CUS + CCPA (0 mg/kg) and Control + Vehicle group (*t* = 5.209, *P* < 0.0001 in distance moved; *t* = 4.924, *P* < 0.001 in center time). One-way ANOVA followed by *post hoc* Bonferroni’s test were performed to compare CUS groups that were given CCPA (0 mg/kg, 0.002 mg/kg, 0.02 mg/kg and 0.2 mg/kg) (*F*_3,28_ = 7.373, *P* < 0.001 in distance moved; *F*_3,28_ = 6.208, *P* < 0.01 in center time). ^#^ *P* < 0.05, ^##^ *P* < 0.01, ^###^ *P* < 0.001, ^####^ *P*< 0.0001, vs. CUS + CCPA (0 mg/kg) group. **d.** Elevated plus maze test (EPM) at 48 h after a single intraperitoneal injection of vehicle or CCPA. Unpaired *t* tests were performed to compare CUS + CCPA (0 mg/kg) and Control + Vehicle group (*t* = 7.903, *P* < 0.0001 in time at open arms; *t* = 2.611, *P* < 0.05 in entrance to open arms), it showed that both time at open arms and entrance to open arms were significantly decreased in the CUS + CCPA (0 mg/kg) group compared with Control + Vehicle group. One-way ANOVA followed by *post hoc* Bonferroni’s test were performed to compare CUS groups that were given CCPA (0 mg/kg, 0.002 mg/kg, 0.02 mg/kg and 0.2 mg/kg) (*F*_3, 36_ = 16.20, *P* < 0.0001 in time at open arms; *F*_3, 36_ = 1.724, *P* > 0.05 in entrance to open arms). ^#^ *P* < 0.05, ^##^ *P* < 0.01, ^###^ *P* < 0.001, ^####^ *P*< 0.0001, vs. CUS + CCPA (0 mg/kg) group. *n* = 8-10 per group. Data were presented as mean ± SEM.

**Fig. S4** **Knockout of CRE (TGACGTCA) sequence on the promotor of rat *Per1* or *Per2* locus by CRISPR/Cas 9.**

**a.** Knockout of CRE sequence (Del_TGACGTCA) on the promotor of *Per1* locus by CRISPR/Cas 9-mediated genomic engineering. **b.** Knockout of CRE sequence (Del_TGACGTCA) on the promotor of *Per2* locus by CRISPR/Cas 9-mediated genomic engineering. **c.** Sanger sequencing verified knockout CRE sequence on the promotor of *Per1* (taking no.60 as an example). **d.** Sanger sequencing verified knockout CRE sequence on the promotor of *Per2* (taking no.6 as an example). WT, wild type. gRNA, guide RNA.
